# Risk evaluation of newly emerging flu viruses based on genomic sequences and AI

**DOI:** 10.1101/2025.04.18.649608

**Authors:** Huiru Li, Yang Feng, Congyu Lu, Ping Fu, Xinglian Wang, Lei Yang, Yuelong Shu, Taijiao Jiang, Dayan Wang, Yousong Peng

## Abstract

The recent resurgence of highly pathogenic avian influenza H5N1 viruses in North America and Europe has heightened global concerns regarding potential influenza pandemics. Despite significant progress in the surveillance and prevention of emerging influenza viruses, effective tools for rapid and accurate risk assessment remain limited. Here, we present FluRisk, an innovative computational framework that integrates viral genomic data with artificial intelligence (AI) to enable rapid and comprehensive risk evaluation of emerging influenza strains. FluRisk incorporates a curated database of over 1,000 experimentally validated molecular markers linked to key viral phenotypes, including mammalian adaptation, mammalian virulence, mammalian transmission, human receptor-binding preference, and antiviral drug resistance. Leveraging these markers, we developed three state-of-the-art machine learning models to predict human adaptation, mammalian virulence, and human receptor-binding potential, all of which demonstrated superior performance compared to traditional approaches such as BLAST, prior models, and baseline classifiers. In addition, a reference-based method was implemented to provide preliminary estimates of human transmissibility and resistance to six commonly used antiviral drugs. To facilitate broad accessibility and practical application, we developed a user-friendly web server that integrates both the molecular marker atlas and predictive tools for influenza virus phenotyping (available at: http://www.computationalbiology.cn/FluRisk/#/). This computational platform offers a valuable resource for the timely risk assessment of emerging influenza viruses and supports global influenza surveillance efforts.

## Introduction

The influenza virus is a negative-sense, single-stranded RNA virus with segmented genomes, classified into four types: A, B, C, and D^1^. Among these, influenza A viruses (IAVs) exhibit the broadest host range, infecting diverse avian species as well as mammals, including humans^2^. IAVs are further categorized into subtypes—such as H1N1, H3N2, and H7N9—based on their surface glycoproteins, hemagglutinin (HA) and neuraminidase (NA)^3^. Over the past two decades, emerging IAVs have posed persistent threats to global public health, exemplified by outbreaks of H7N9 and H5N1^4–7^. Notably, the highly pathogenic avian influenza (HPAI) H5N1 virus has demonstrated alarming cross-species transmission, infecting over 50 mammalian species such as cattle, sheep, pig, cats, etc^8–12^. This expanding host range has intensified global concerns about its pandemic potential. Effective prevention and control of emerging IAVs remain a critical priority for public health worldwide.

Risk assessment of emerging influenza viruses is a prerequisite for effective pandemic control. Previously, the European Food Safety Authority (EFSA) proposed a risk assessment framework to rank animal influenza strains according to their potential to infect humans in the FLURISK project^13^. The American CDC has developed the Influenza Risk Assessment Tool (IRAT) to assess the potential pandemic risk posed by influenza A viruses that are not currently circulating in people^14^. The World Health Organization (WHO) also developed the Tool for Influenza Pandemic Risk Assessment (TIPRA) to evaluate the public health threat posed by emerging influenza strains^15^. This framework integrates virological, epidemiological, and clinical data to systematically assess both the likelihood of emergence and potential impact of pandemic influenza viruses^15^. However, obtaining virological information such as host, virulence and transmission ability for newly emerging viruses remains a significant challenge, as it often requires extensive laboratory testing and epidemiological studies. Such inherent methodological complexities, coupled with the protracted timelines of conventional studies, often result in delayed availability of pivotal evidence. Consequently, this temporal gap may compromise timely and accurate pandemic risk assessments during early emergence events.

The development of computational methods for rapid viral phenotype prediction could significantly enhance influenza risk assessment, particularly given the increasing availability of influenza genomic data and advances in artificial intelligence (AI), including deep learning and large language models. As of April 15, 2025, the GISAID database contains over 3 million genomic sequences from 637,120 influenza virus strains^16^, providing a rich resource for data-driven modeling. Numerous AI-based methods have been developed to infer key viral phenotypes such as host specificity, virulence, and antigenic variation, directly from genomic sequences^17–19^. For instance, we developed the PREDAC method based on the Naïve Bayes algorithm to predict influenza antigenic variation from hemagglutinin (HA) protein sequences^17^; Yin et al. developed a computational framework integrating prior viral knowledge to predict mammalian virulence from influenza genomic data^18^. Borkenhagen et al. systematically reviewed machine learning (ML) approaches for influenza phenotype prediction, highlighting the growing role of AI in virology^20^. Despite these advances, the integration of such models into routine influenza surveillance and risk assessment remains limited. Key barriers include suboptimal model performance in real-world settings and the lack of user-friendly, deployable tools for public health applications.

Molecular markers, defined as specific amino acid residues at defined positions that correlate with distinct influenza virus phenotypes, provide a potential framework for rapid phenotypic inference to support risk assessment. The PB2-627K, for instance, is a well-characterized marker associated with enhanced mammalian pathogenicity ^21,22^, while numerous other markers have been experimentally validated for phenotypes including host adaptation, virulence, antigenic variation, drug resistance, and human transmissibility^23–27^. These markers offer a potential shortcut for roughly inferring viral phenotypes, thereby facilitating rapid risk assessments in influenza virus surveillance. However, their practical utility faces two key limitations: first, most markers were identified using limited viral strains, and their generalizability across diverse genetic backgrounds remains uncertain due to epistatic interactions between amino acid sites; second, the lack of a unified, curated database results in fragmented knowledge, with experimentally validated markers dispersed across the literature rather than systematically organized for operational use.

Building upon our previous development of FluPhenotype - a one-stop platform for predicting influenza A virus host range and antigenicity^28^ - we now introduce FluRisk, an advanced AI-driven platform designed to enable rapid and comprehensive risk assessment of emerging influenza viruses through genomic sequence analysis. FluRisk integrates two key innovations: (1) a systematically curated, up-to-date atlas of influenza molecular markers, and (2) machine learning models capable of predicting five critical phenotypic traits (human adaptation, mammalian virulence, human receptor-binding affinity, transmissibility, and drug resistance). To facilitate practical implementation, we have deployed FluRisk as an accessible web server, providing researchers and public health professionals with a powerful tool for real-time influenza risk evaluation and surveillance.

## Results

### Overview of FluRisk

Figure 1A provides an overview of the FluRisk platform architecture. The system accepts influenza virus genomic or protein sequences as sole input requirements. FluRisk incorporates a comprehensive database of experimentally validated molecular markers associated with five critical phenotypic categories: (1) mammalian adaptation, (2) mammalian virulence, (3) human receptor-binding preference, (4) mammalian transmissibility, and (5) resistance profiles against six clinically relevant anti-influenza drugs. The platform automatically extracts and analyzes relevant molecular markers from submitted sequences through its integrated processing pipeline. Multiple computational methods were developed that leverage both these molecular markers and direct sequence features to predict phenotypic characteristics and antigenic properties. FluRisk generates two primary outputs: (1) a detailed report of identified phenotype-associated molecular markers, and (2) quantitative risk scores for each predicted phenotype, enabling early warning assessment of emerging viral threats. Technical details regarding the molecular marker database construction and computational prediction methodologies are presented in subsequent sections.

**Figure 1.**
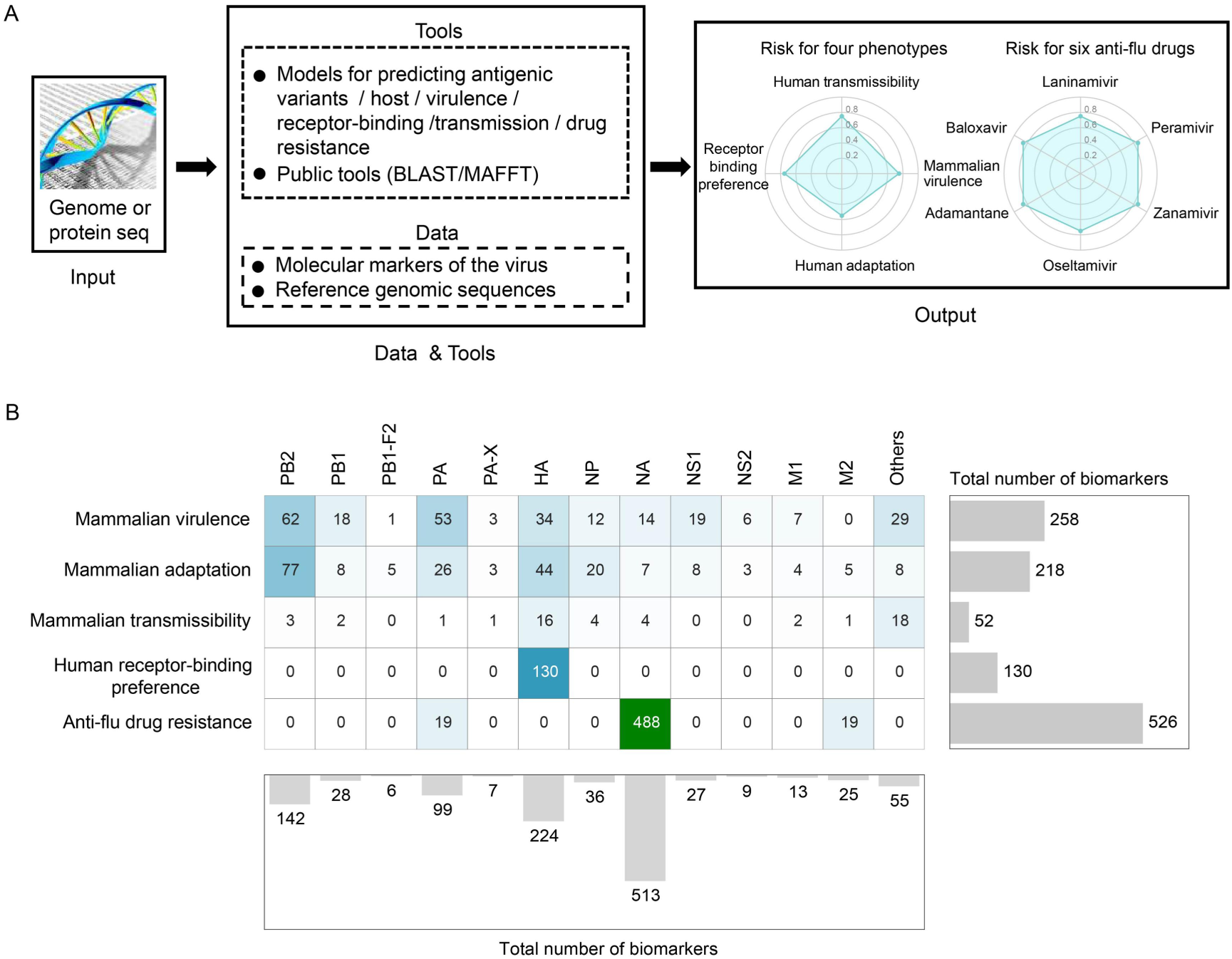
Overview of FluRisk (A) and Summary of molecular markers of important influenza virus phenotypes (B)

### Summary of molecular markers associated with flu phenotypes

FluRisk integrates a comprehensive database of 1,184 experimentally validated molecular markers associated with influenza phenotypes, which were manually curated from published literature (see Methods). These molecular markers cover five major phenotypic categories. For mammalian adaptation, we identified 218 molecular markers distributed across all viral proteins, with PB2 containing the highest number (77 markers, including well-characterized markers like 627K and 701N), followed by HA (44 markers), PA (26 markers), and NP (20 markers), while other proteins each contained fewer than 10 markers. Regarding mammalian virulence, the database includes 258 molecular markers present on all viral proteins except M2, with PB2 again showing the highest number (62 markers), followed by PA (53 markers), HA (34 markers), NS1 (19 markers) and PB1 (18 markers). For human receptor-binding preference, we curated 130 molecular markers specifically located on the HA protein across 7 HA subtypes. The mammalian transmission phenotype was associated with 52 molecular markers distributed across multiple proteins, with HA containing about one-third of these markers, and interestingly, one-third of markers were located on two or more proteins, such as the combination of H3-225G and N2-315N. For drug resistance, we collected molecular markers for six anti-influenza drugs: oseltamivir (168 markers), zanamivir (155 markers), peramivir (106 markers), laninamivir (58 markers), baloxavir (20 markers), and adamantane (19 markers), which were classified into four resistance levels (low, moderate, high, and unknown) based on IC50 values. Among these, 488 markers were located on the NA protein of 3 NA subtypes, while the remaining 38 markers were found on PA and M2 proteins.

Protein-level analysis revealed distinct distributions of molecular markers across viral proteins. The NA protein harbored the highest number of molecular markers (n=513), with the vast majority (95.1%) associated with drug resistance phenotypes. Following NA, the HA protein contained the second largest marker collection (n=224), of which 58% were linked to human receptor-binding preference. The PB2 protein ranked third in marker abundance (n=142), with its markers predominantly involved in mammalian adaptation (54.2%) and mammalian virulence (43.7%) phenotypes. Additionally, the PA protein contained a substantial number of markers (n=99) distributed across three phenotypic categories: mammalian virulence (53.5%), mammalian adaptation (26.3%), and drug resistance (19.2%). This protein-specific marker distribution reflects their respective functional roles in influenza virus biology and pathogenesis.

### Machine learning-based prediction of influenza virus host

Then we built several computational methods to predict the influenza virus phenotypes based on molecular markers. For host prediction, we first analyzed the distribution of mammalian-adaptation markers across viral genomes. Comparative analysis revealed a striking disparity: human influenza viruses carried significantly more mammalian-adaptation markers than avian strains (median: 84 *vs.* 18; Figure 2A). Protein-level examination showed this pattern held across nearly all viral proteins (Figure 2B). Among these, PB2 exhibited the most pronounced difference, with human virus strains containing a median of 40 markers compared to avian strains. The NP and PA proteins followed, with median counts of 17 and 12 markers respectively in human influenza viruses. These quantitative differences in marker distribution formed the basis for our host-prediction algorithm.

**Figure 2.**
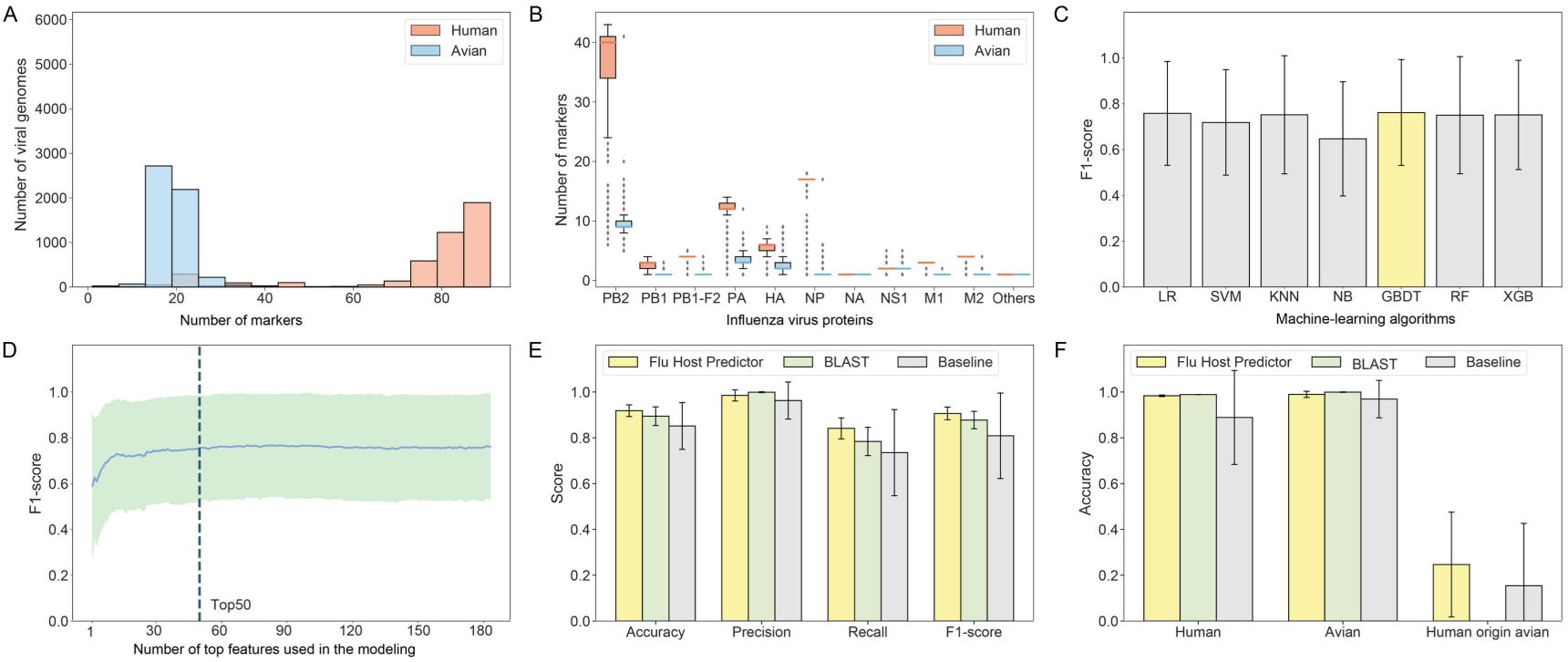
Host prediction of influenza viruses using mammalian-adaptation molecular markers. (A) Genome-wide comparison of mammalian-adaptation marker counts between human and avian influenza viruses. (B) Protein-specific distribution of mammalian-adaptation markers across viral proteins. (C) Machine learning algorithm selection for Flu Host Predictor (FHP) based on validation set performance. (D) Feature selection optimization for FHP model construction. (E) Comparative performance evaluation of FHP, BLAST, and baseline methods on independent test data. (F) Host-specific accuracy analysis of all three prediction methods across different influenza virus categories.

Building upon these findings, we developed Flu Host Predictor (FHP), a machine learning model for classifying human versus avian influenza viruses based on mammalian-adaptation molecular marker profiles. The model construction workflow is detailed in Figure S1 and the Methods section. Comparative evaluation of multiple algorithms including Logistic Regression (LR), Support Vector Machine (SVM), k-Nearest Neighbors (KNN), Naive Bayes (NB), Random Forest (RF), XGBoost (XGB), and Gradient Boosting Decision Tree (GBDT) revealed GBDT as the optimal choice, achieving the highest F1-score (0.762) when using all features (Figure 2C). To optimize model performance while minimizing overfitting, we selected the top 50 most important features (Figure 2D). The final model demonstrated robust performance on the testing dataset, with an average F1-score of 0.906, along with accuracy (0.918), precision (0.985), and recall (0.841) metrics (Figure 2E). Benchmarking against conventional approaches showed that while BLAST (using best-hit host assignment) and a baseline method (counting total mammalian-adaptation markers) (see Methods) achieved comparable precision (∼100%), their recall rates were substantially lower than FHP. Host-specific performance analysis revealed high accuracy for conventional human and avian influenza viruses across all methods, but notably poor performance for human-origin avian influenza viruses (average accuracy of 0.247, 0 and 0.154 for FHP, BLAST and baseline method, respectively) (Figure 2F).

### Machine learning-based prediction of mammalian virulence in influenza viruses

We next developed a machine learning approach to predict mammalian virulence using molecular markers associated with this phenotype. The analysis began with the compilation of a carefully curated dataset comprising 659 influenza virus strains with experimentally validated virulence phenotypes, obtained from published literature. This dataset included 198 high-virulence and 461 low-virulence strains from 21 subtypes (Figure 3A). The H5N1 subtype was most prevalent in our collection, with 101 high-virulence and 143 low-virulence strains, followed by H1N1 (41 high-virulence and 120 low-virulence strains). All other subtypes had less than 100 viral strains. Comparative analysis revealed that the high-virulence strains had larger number of molecular markers than the low-virulence strains (p-value = 6.60e-04 in the Wilcoxon rank-sum test) (Figure 3B). The former had a median of 45 molecular markers, while the latter had a median of 43 molecular markers. Protein-specific examination demonstrated that this difference was particularly pronounced in the PB2 and HA proteins, where high-virulence strains exhibited more markers than their low-virulence counterparts (Figure 3C).

**Figure 3.**
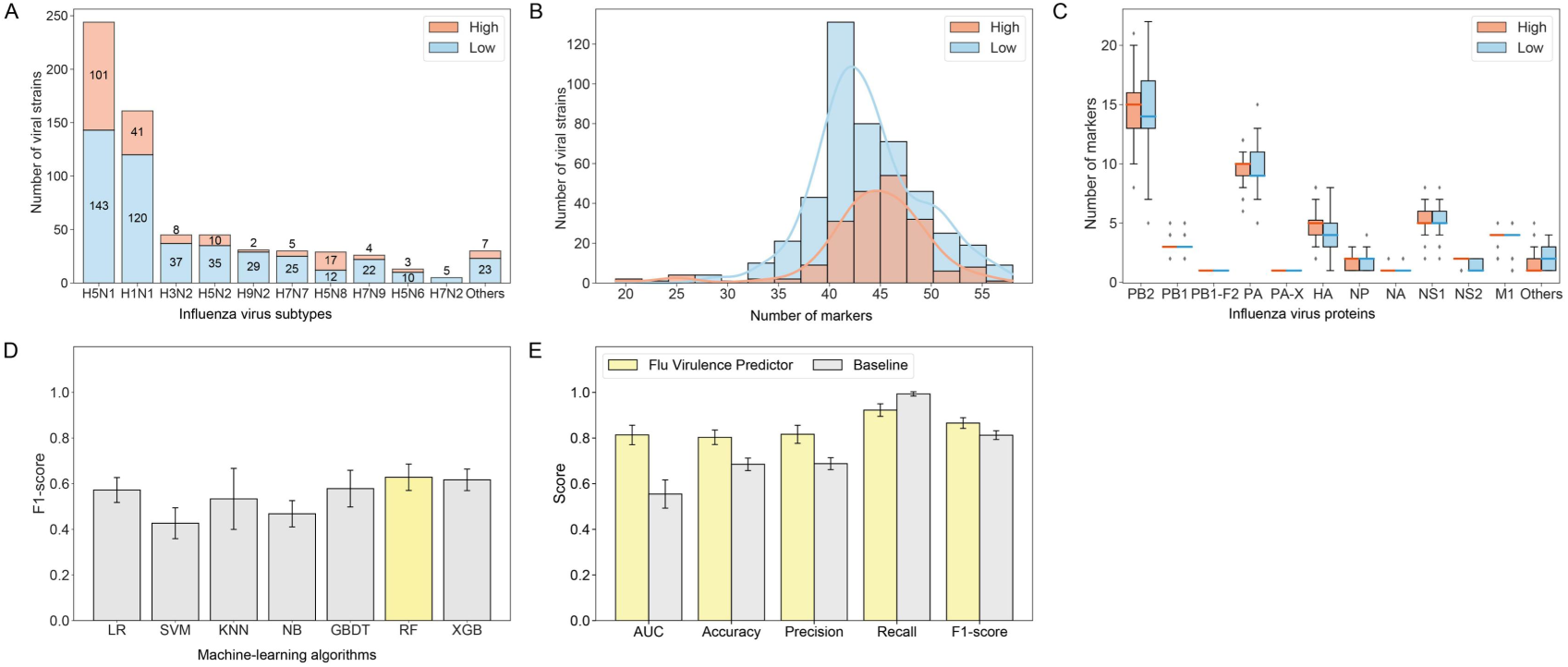
Machine learning-based prediction of influenza virus mammalian virulence. (A) Distribution of high-virulence and low-virulence influenza virus strains across different subtypes. (B) Genome-wide comparison of mammalian-virulence-associated molecular markers between high- and low-virulence strains. (C) Protein-specific distribution of virulence-associated molecular markers in high- and low-virulence strains. (D) Algorithm selection for Flu Virulence Predictor (FVP) based on F1 scores across multiple machine learning approaches. (E) Comparative performance evaluation of FVP versus baseline method on independent test dataset, reporting accuracy, precision, recall, F1-score and AUC metrics.

Building on these analysis, we developed Flu Virulence Predictor (FVP), a computational method for predicting mammalian virulence of influenza viruses using virulence-associated molecular markers. Following a workflow analogous to FHP, we evaluated multiple machine learning algorithms (LR, SVM, KNN, NB, GBDT, XGB, and RF) and selected random forest (RF) as the optimal approach based on F1-score performance. To mitigate overfitting, we implemented feature selection using the VSURF method (see Methods). Evaluation of FVP on the testing dataset showed that FVP had an overall F1-score and AUC of 0.866 and 0.814, respectively. Specifically, the accuracy, recall and precision for FVP were 0.803, 0.922 and 0.816, respectively. Comparative analysis against a baseline method (which classified virulence solely based on total molecular marker count) showed FVP’s superior performance across most metrics (Figure 3E). While the baseline method achieved higher recall (0.993 vs 0.922), FVP showed substantial improvements in accuracy (0.803 vs 0.685), precision (0.816 vs 0.688), F1-score (0.866 vs 0.812), and AUC(0.814 ys 0.555).

To further validate FVP’s predictive capability, we conducted benchmarking against two established virulence prediction methods developed by Yin et al. - ViPal^18^ and VirPreNet ^29^ - using their original dataset. This comparative analysis was necessary as we were unable to fully reproduce their methods due to unavailability of certain critical parameters. ViPal employs a generalized machine learning framework incorporating genomic sequences along with prior viral mutation and reassortment data, while VirPreNet utilizes a weighted ensemble convolutional neural network with ProtVec encoding for virulence prediction. As detailed in Table S1, FVP demonstrated superior performance across all evaluation metrics except recall, where it achieved comparable results to VirPreNet.

### Machine learning approach for predicting receptor-binding preference of influenza viruses

Then, we continued to develop a computational method to predict influenza virus receptor-binding specificity using machine learning algorithms. Our analysis began with the compilation of a carefully curated dataset comprising 620 influenza A virus strains with experimentally validated receptor-binding preferences (human vs avian), sourced from public databases and literature. The receptor-binding data of more than half of viral strains (58.1%) were obtained using the glycan array; 39.0% of viral strains were determined using ELISA; the remaining viral strains were determined using BLI (1.9%) and SPR (1.0%) (Figure 4A). The dataset contains 321 and 299 influenza virus strains with human (α2-6-linked) and avian-receptor (α2-3-linked) binding preferences, respectively, from 33 influenza A virus subtypes. The subtype distribution showed H1N1 as the most prevalent (103 human-binding and 64 avian-binding strains), followed by H3N2 and H9N2 subtypes, each containing over 100 strains. This comprehensive dataset provided the foundation for our predictive modeling approach.

**Figure 4.**
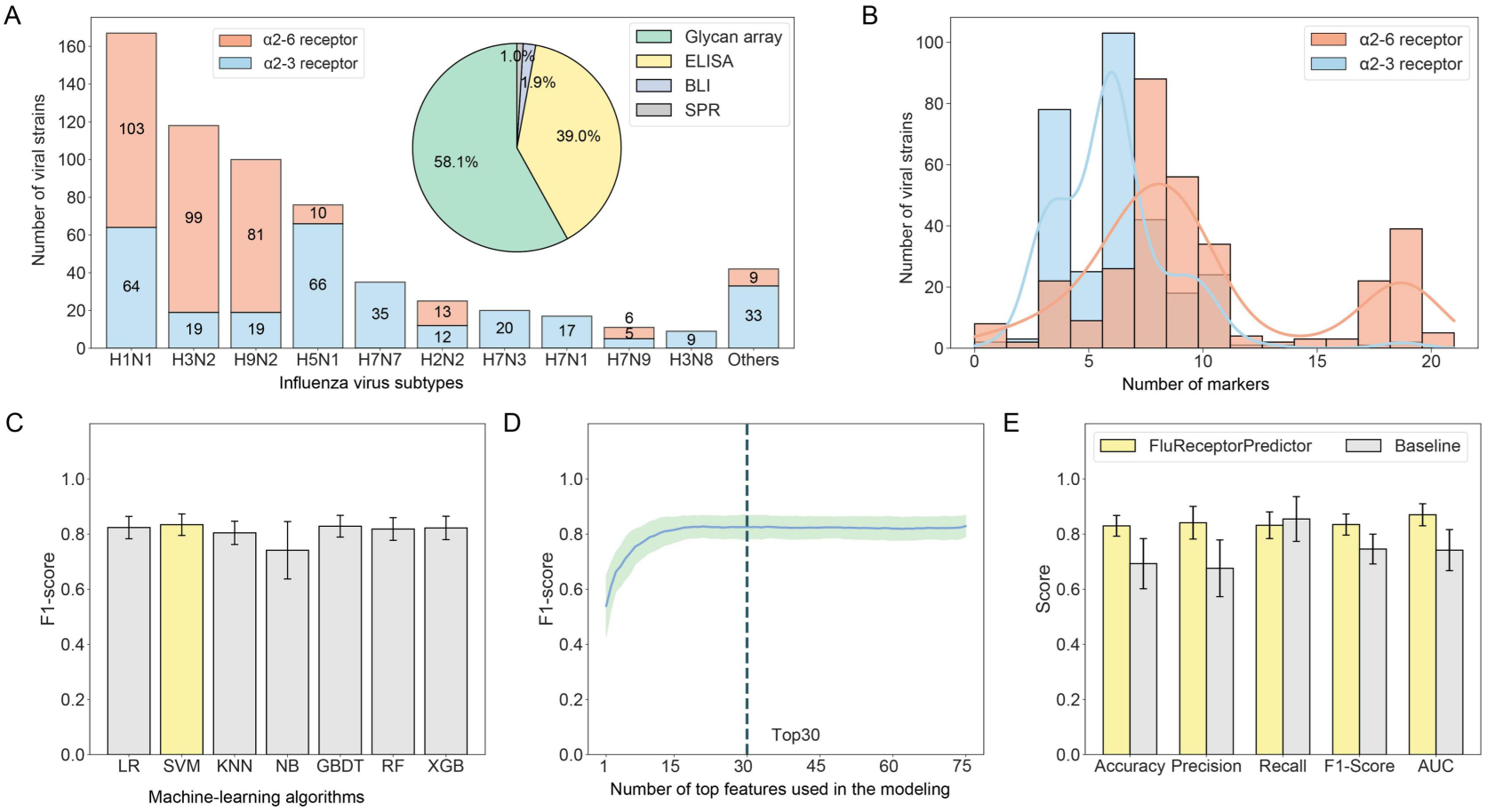
Machine learning prediction of influenza virus receptor-binding preference using molecular markers. (A) Distribution of viral strains with human (α2-6-linked sialic acid) versus avian (α2-3-linked) receptor preference across subtypes. Inset pie chart shows proportion of experimental methods used for receptor-binding determination (glycan array, ELISA, BLI, SPR). (B) Comparative analysis of human-receptor-binding-associated molecular markers between human-preferring and avian-preferring strains. (C) Algorithm selection for Flu Receptor-binding preference Predictor (FRP) based on F1-score performance across multiple machine learning algorithms. (D) Feature selection optimization identifying the top 30 molecular markers. (E) Performance evaluation of FRP versus baseline method on independent test dataset, reporting accuracy, precision, recall, F1-score, and AUC metrics.

Comparative analysis of molecular markers associated with human-receptor-binding preferences revealed a highly significant difference in marker numbers between strains with human versus avian receptor-binding preferences (p-value = 3.42e-31 in the Wilcoxon rank-sum test). The former had a median of 9 markers, compared to 6 markers in the latter (Figure 4B). Leveraging these findings, we developed the Flu Receptor-binding preference Predictor (FRP) using a workflow analogous to FHP (Figure S1). Algorithm selection demonstrated that support vector machine (SVM) achieved the highest F1-score among evaluated algorithms (LR, KNN, NB, GBDT, RF, and XGB) (Figure 4C). Feature selection identified the top 30 most important markers to optimize model performance while minimizing overfitting (Figure 4D). When tested on the testing dataset, FRP had a F1-score of 0.834 and AUC of 0.870 (Figure 4E). Specifically, the average accuracy, precision and recall was 0.830, 0.841 and 0.832, respectively. Benchmarking against a baseline method (which classified receptor-binding preference solely based on molecular marker count) showed FRP’s superior performance across all metrics except recall (Figure 4E).

### Reference-based estimation of human transmission potential and drug-resistance of influenza viruses

Due to insufficient data for machine learning model development, we established a reference-based approach to estimate influenza virus human transmission potential. We first constructed a reference library comprising 500 representative human and 500 avian influenza virus strains (see Methods). Comparative analysis revealed a highly significant difference (p-value=3.16e-135 in Wilcoxon rank-sum test) in mammalian-transmission-associated molecular marker number between human (median=5 markers) and avian (median=2 markers) strains (Figure 5A, left panel), suggesting marker quantity correlates with human transmission potential. Then, we generated a cumulative distribution of mammalian-transmission markers across all reference strains (Figure 5A, right panel). For a given virus, we: (1) quantified its mammalian-transmission markers in the genome, (2) determined its percentile rank in the reference distribution, and (3) used this normalized ranking ratio (illustrated by red dashed line in Figure 5A) as its estimated human transmission ability score. This approach provides a standardized, quantitative measure of transmission potential relative to known human and avian influenza viruses.

**Figure 5.**
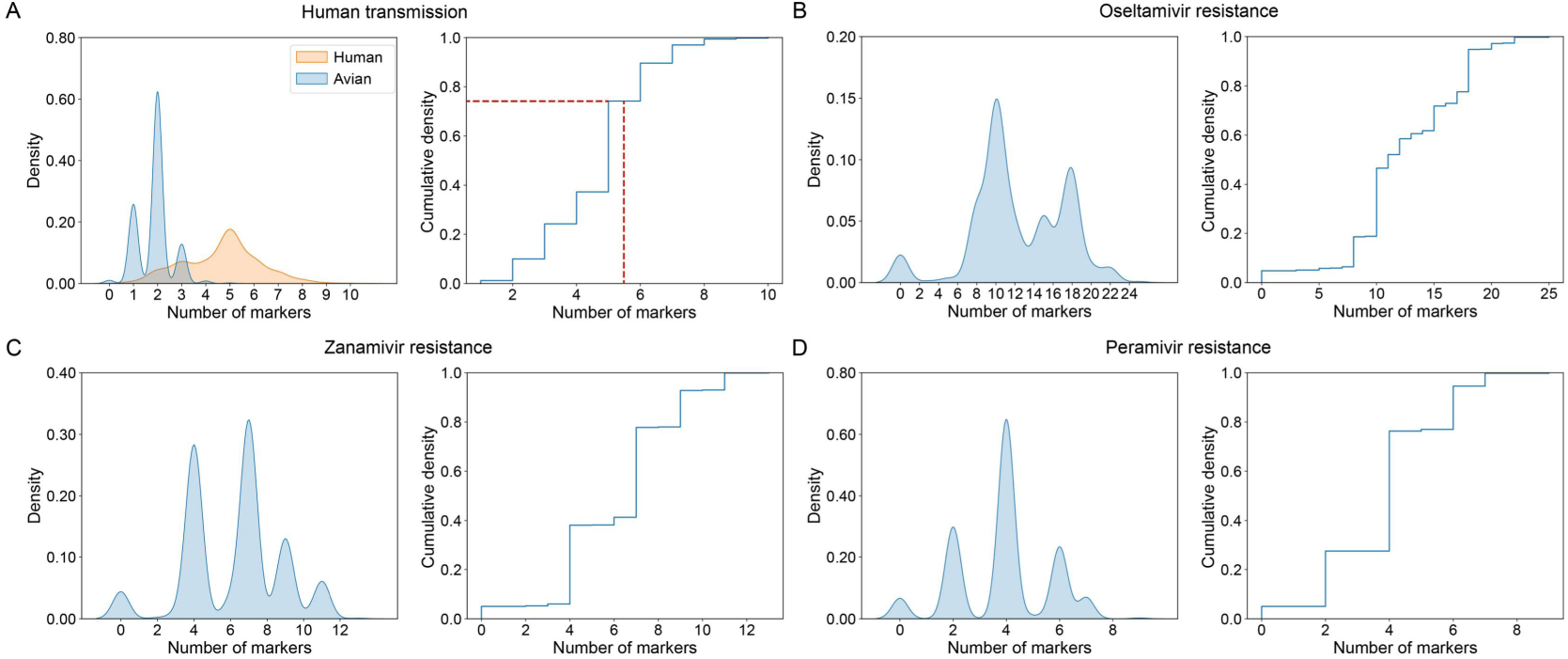
Estimation of human transmission and drug-resistance of influenza viruses based on molecular markers using a reference-based method. (A) The left panel shows the comparison of the number of molecular markers associated with mammalian transmission in reference human (orange) and avian (blue) influenza virus strains, while the right panel shows the cumulative distribution of the number of molecular-markers associated with mammalian transmission in virus genomes using all reference strains. The red dashed line shows an example of estimating the ability of human transmission for a given virus. (B)-(D) shows the distribution of the number of molecular markers and the cumulative distribution of the number of molecular markers associated with Oseltamivir, Zanamivir, and Peramivir, respectively.

We extended our reference-based approach to estimate drug resistance profiles for influenza viruses. A curated collection of 1,000 representative human influenza virus strains served as the reference dataset. For each of the six clinically relevant anti-influenza drugs, we: (1) quantified resistance-associated molecular markers in all reference strains, and (2) constructed individual cumulative distribution. Analysis revealed distinct resistance marker patterns across drug classes (Figures 5B-D & Figure S2). The reference strains exhibited a median of 11 oseltamivir-resistance markers (Figure 5B), while strains carried less then 10 resistance markers for other drugs such as zanamivir (Figure 5C) and peramivir (Figure 5D).

### Development of a web server for FluRisk

To facilitate public access to our database and computational tools, we developed the user-friendly FluRisk web server (http://www.computationalbiology.cn/FluRisk/#/) with an intuitive interface (Figure 6A). Besides the tools mentioned above, the antigenicity of influenza viruses was predicted using the PREDAC method that was developed in our previous studies. The platform’s core functionality provides comprehensive risk assessment for emerging influenza viruses by analyzing input genomic or proteomic sequences to identify phenotype-associated molecular markers and generate quantitative risk scores for five critical phenotypes (Figure 6B). The server hosts a searchable molecular marker database organized by phenotype or protein, with detailed annotations covering functional consequences, supporting literature, and protein structure visualization, while also allowing community contributions through a submission interface for newly discovered markers. Supplementary utilities include HA/NA subtype-specific amino acid numbering conversion and antigenic variant prediction from protein sequences. Designed with modular architecture, FluRisk enables continuous integration of new prediction models and database updates to support ongoing research and surveillance efforts.

**Figure 6.**
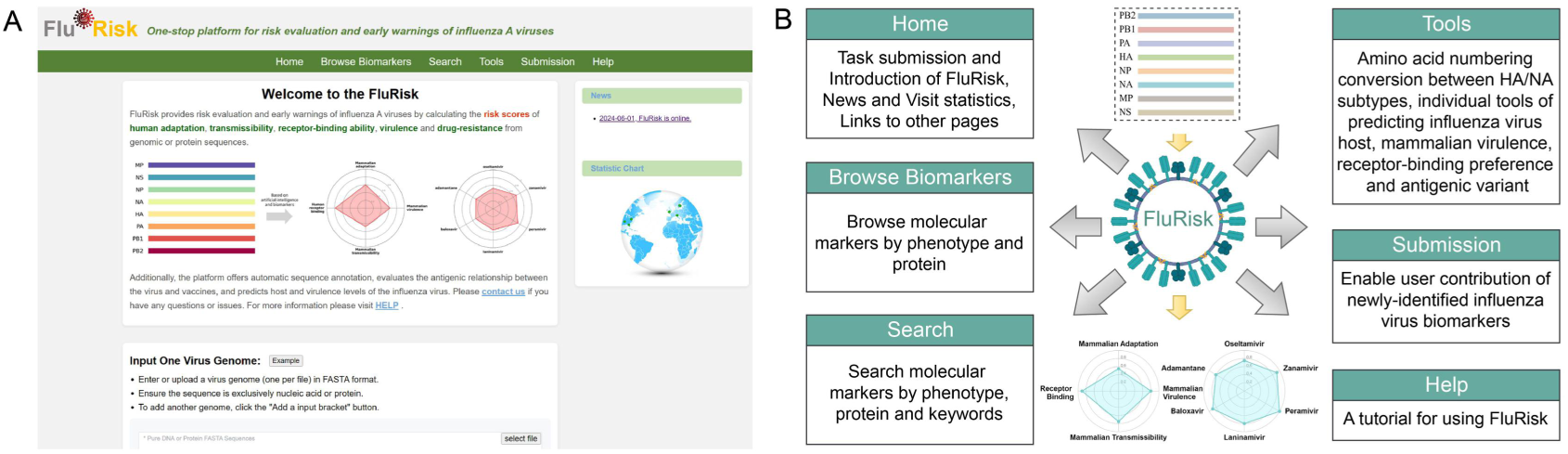
Construction of the web server of FluRisk. (A) The homepage of the FluRisk web server. (B) The structure of the FluRisk web server.

## Discussions

This study established a comprehensive atlas of molecular markers associated with five critical influenza virus phenotypes, serving as both a valuable resource for viral variant annotation and a foundation for rapid phenotypic inference in emerging strains. Leveraging this marker database, we developed three state-of-the-art machine learning models (for host prediction, mammalian virulence assessment, and receptor-binding preference determination) and a novel reference-based method (for estimating human transmissibility and drug resistance profiles) - all utilizing genomic sequence data as input. These computational approaches enable timely phenotypic predictions that significantly enhance risk assessment capabilities for emerging influenza threats. The integration of these predictive tools with our molecular marker database in the publicly accessible FluRisk web server creates a powerful platform for early warning and surveillance of potentially dangerous viral variants in real-world public health practice.

The WHO’s TIPRA framework provides a comprehensive system for influenza pandemic risk assessment by integrating virological, epidemiological, and clinical data ^15^. However, its practical implementation faces significant challenges when evaluating emerging viruses, as critical phenotypic data are often unavailable during early outbreaks. Conventional determination of viral biological properties requires labor-intensive experimental characterization, which becomes particularly problematic for dangerous emerging strains requiring high-containment biosafety facilities. These constraints inevitably delay risk assessment and hinder large-scale surveillance of emerging variants. In contrast, computational approaches leveraging viral genomic sequences offer a rapid and cost-effective alternative, especially given the increasing accessibility of viral genome sequencing. While several genomic prediction methods have been developed (as comprehensively reviewed by Borkenhagen et al ^20^.), our study advances the field in two key aspects: First, our machine learning models incorporate experimentally validated molecular markers as predictive features, providing more biologically interpretable and potentially more reliable predictions compared to previous sequence-based approaches. Second, we have compiled substantially larger benchmark datasets for mammalian virulence and receptor-binding preference prediction than those used in prior studies, establishing valuable resources for future methodological development in this field.

The phenotypic characteristics of influenza viruses, including virulence, host adaptation, and human transmissibility, are polygenic traits resulting from complex genotype-phenotype relationships^30,31^. Deciphering these relationships requires both sophisticated computational approaches and extensive datasets linking viral genomes to phenotypic outcomes. However, current phenotypic data for influenza viruses remain limited in scale - for instance, our mammalian virulence and receptor-binding models incorporated fewer than 1,000 viral strains. This scarcity highlights two critical needs: (1) high-throughput experimental methods for phenotypic characterization, and (2) advanced computational approaches to extract phenotypic information from genomic data, given the predominant role of viral genomes in determining these traits. Recent advances in large language models (LLMs) have demonstrated superior performance over traditional machine learning in predicting various viral characteristics^32–34^. The E2VD framework developed by Nie et al. ^34^, for example, effectively predicts mutation effects on SARS-CoV-2 binding affinity, expression, and antibody escape using a pretrained language model. However, these models have been built mainly based on the deep mutational scanning (DMS) data. They exhibit significant performance degradation when applied to viral strains with genetic backgrounds divergent from their training data (e.g., different SARS-CoV-2 variants) ^34^. This limitation poses particular challenges for influenza research, where emerging strains may belong to diverse subtypes (H5N1, H7N9, H10N8, etc.) with substantially different genetic contexts. While current DMS-derived LLMs may not be optimal for influenza phenotype prediction, the expanding availability of DMS data across diverse influenza strains may eventually enable development of specialized LLMs capable of accurately modeling the complex genome-phenotype relationships in influenza viruses.

While this study provides valuable resources and tools for influenza risk assessment, several limitations should be acknowledged. First, although we have compiled an extensive atlas of phenotype-associated molecular markers, it remains inherently incomplete due to the continuous emergence of novel viral mutations. To address this, we have implemented a mechanism for continuous database updates through community contributions via the FluRisk web server. Second, while our computational models demonstrate superior performance compared to existing methods, there remains room for improvement in prediction accuracy. Future integration of more advanced algorithms, such as large language models, coupled with expanded phenotypic datasets, may enhance model performance. Third, our current risk evaluation framework relies solely on genomic data. Incorporating additional epidemiological, clinical, and ecological parameters - as employed in WHO’s comprehensive assessment approach - could significantly improve risk evaluation for emerging influenza viruses. These limitations highlight important directions for future research and development in this field.

## Conclusion

This study makes three significant contributions to influenza research and surveillance: (1) establishment of a comprehensive, curated atlas of molecular markers associated with five critical influenza virus phenotypes, serving as an invaluable resource for the scientific community; (2) development of a suite of advanced computational methods that enable rapid phenotypic prediction directly from viral genomic sequences, overcoming the limitations of traditional experimental approaches; and (3) construction of a user-friendly FluRisk web server, integrating both predictive tools and molecular marker data into a unified platform for early detection and risk assessment of emerging influenza viruses. Together, these advances provide researchers and public health professionals with powerful new capabilities for influenza surveillance and pandemic preparedness.

## Methods

### Compilation of phenotype-associated molecular markers

The molecular markers (defined as specific amino acid residues at particular positions that correlate with phenotypic changes) were systematically collected from two primary sources: (1) 397 previously curated markers from our FluPhenotype platform^28^, and (2) an exhaustive literature review of PubMed entries published between January 1, 2019 and December 31, 2022 since FluPhenotype has collected molecular markers from public literatures published before 2019. The literature collection process involved multiple filtering steps: initial screening identified 8,238 influenza-related publications, which were narrowed to 1,431 papers containing mutation data using a custom Python script. Each publication underwent manual review to identify markers associated with five key phenotypes: mammalian adaptation, mammalian virulence, human receptor-binding specificity, mammalian transmissibility, and resistance to six antiviral drugs (oseltamivir, zanamivir, peramivir, laninamivir, baloxavir marboxil, and amantadine). For drug resistance markers, we recorded available IC50 values and classified them according to WHO guidelines based on fold-change in IC50 (IC50-FC): low resistance (2 < IC50-FC ≤ 10), moderate resistance (10 < IC50-FC ≤ 100), high resistance (IC50-FC > 100), and unknown resistance (when IC50 data were unavailable). Following integration of both data sources, we established a comprehensive collection of 1,184 molecular markers, which have been made publicly accessible through the FluRisk web server (http://www.computationalbiology.cn/FluRisk/#/).

### Influenza virus genomic sequence data collection and processing

Viral genomic sequences were retrieved from the GISAID database on May 20, 2023. Initial quality filtering retained only complete genomes, yielding 18,966 genomes. We subsequently applied CD-HIT (version 4.8.1)^35^ to reduce sequence redundancy at a 99.9% similarity threshold, resulting in three distinct datasets: 5,257 avian influenza virus genome sequences, 4,155 human influenza virus genome sequences, and 349 human-origin avian influenza virus genome sequences. These non-redundant sequences served as the foundation for developing our host prediction models.

For specific analytical applications, we generated two reference sequence subsets through random sampling: (1) 1,000 human influenza virus genomes for drug resistance risk assessment, and (2) balanced sets of 500 human and 500 avian influenza virus genomes for human transmission risk evaluation.

### Compilation of influenza virus mammalian virulence data

The mammalian virulence dataset was systematically curated by integrating data from two comprehensive studies by Yin et al.^18^ and Peng et al.^36^. To ensure consistent virulence classification across studies, we established standardized criteria: viral strains demonstrating either (1) high mortality at low infection doses or (2) a 50% lethal dose (LD50) < 10^3^ were classified as high-virulence, while all others were designated as low-virulence. This rigorous classification yielded a final dataset of 659 characterized viral strains, comprising 198 high-virulence and 461 low-virulence isolates. Corresponding genomic sequences for these strains were obtained through a combination of source publications and the GISAID database.

### Compilation of influenza virus receptor-binding data

Receptor-binding specificity data for influenza viruses were systematically collected from two primary sources. First, we retrieved 606 influenza virus-related records from the Consortium for Functional Glycomics (CFG) database on December 18, 2024, which contained glycan array measurements of viral binding to α2-3-linked (avian-type) and α2-6-linked (human-type) sialic acid receptors^37^. After cross-referencing with hemagglutinin (HA) sequence availability from both CFG and GISAID databases, we obtained 360 viral strains with complete receptor-binding profiles and corresponding HA sequences. Second, we supplemented this dataset with literature-reported binding measurements obtained through three established biophysical techniques: (1) surface plasmon resonance (SPR), (2) biolayer interferometry (BLI) - both quantifying binding affinity through equilibrium dissociation constants (Kd) derived from optical biosensor measurements - and (3) enzyme-linked immunosorbent assay (ELISA), which assesses binding through Kd derived from absorbance values^38–40^. For all methods, stronger binding was indicated by smaller Kd measurements. The combined dataset comprised 620 viral strains with experimentally validated binding profiles for both receptor types. We classified receptor-binding preference based on relative binding affinities: strains demonstrating stronger binding to α2-6-linked glycans were designated as human receptor-preferring, while those with greater affinity for α2-3-linked glycans were classified as avian receptor-preferring.

### Framework for predicting the flu host, virulence and receptor-binding preference using machine learning algorithms

Figure S1 illustrates the comprehensive workflow for predicting influenza host, mammalian virulence, and receptor-binding preference through machine learning analysis of genomic sequences. The initial step involves extracting molecular markers associated with specific phenotypes (e.g., human adaptation) from viral genomic or protein sequences, which serve as predictive features in subsequent modeling. The dataset undergoes systematic partitioning into training/validation (80%) and independent testing (20%) subsets, with the training portion further divided 8:2 for model development and validation purposes. This stratified approach enables algorithm selection, feature optimization, and model training using the training/validation data, while reserving the testing set for unbiased performance evaluation. To account for potential partitioning variability, the entire procedure was iterated 100 times. For host prediction specifically, given the substantial dataset size and diverse HA/NA subtype representation, we implemented subtype-stratified sampling across training, validation, and testing sets to prevent subtype overlap, thereby ensuring rigorous assessment of model generalizability to novel influenza variants.

### Machine-learning algorithms and feature selection for predicting flu virus host, virulence and receptor-binding preference

We evaluated seven established machine-learning algorithms for influenza phenotype prediction: Logistic Regression (LR), Support Vector Machine (SVM), k-Nearest Neighbors (KNN), Naive Bayes (NB), Gradient Boosting Decision Tree (GBDT), Random Forest (RF), and XGBoost (XGB), implemented through scikit-learn (version 3.5.1) ^41^ in Python (version 3.7.12) using their respective classifier functions: SVC for SVM, RandomForestClassifier for RF, LogisticRegression for LR, KNeighborsClassifier for KNN, GaussianNB for NB, GradientBoostingClassifier for GBDT, and XGBClassifier for XGB, with default parameters employed during algorithm selection. For host prediction and receptor-binding preference, GBDT and SVM emerged as the optimal algorithms respectively, with feature selection performed through permutation importance-based forward selection where features were ranked by importance scores from the permutation_importance function applied to the training set, then sequentially incorporated into models while monitoring F1-score improvements on the validation set to identify the optimal feature subset. For mammalian virulence prediction where RF demonstrated superior performance, we employed the VSURF (Variable Selection Using Random Forests) ^42^ method for feature selection, leveraging its inherent compatibility with random forest architectures to identify the most predictive molecular marker subsets.

### BLAST-based host prediction method

To determine the host origin for influenza virus strains in the testing dataset, we first constructed a comprehensive protein sequence library comprising all viral strains from the training and validation datasets, followed by performing BLASTp (version 2.13.0+)^43^ searches against this reference library using each query virus’s protein sequences, with the final host prediction determined through a majority voting mechanism that integrated results across all individual protein matches.

### Baseline prediction method

The baseline prediction approach estimated influenza virus phenotypes (host, mammalian virulence, and receptor-binding preference) solely based on the cumulative count of phenotype-associated molecular markers present in each viral genome, with optimal classification thresholds established through analysis of the training and validation datasets.

### Evaluation criteria for computational methods

We introduced five common performance metrics (i.e. Accuracy, Precision, Recall, F1-Score, and AUC) for method evaluation and comparison. These metrics are defined as

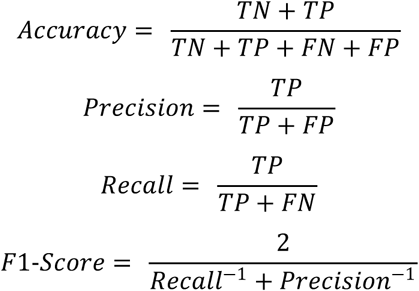

where TP, TN, FP and FN denote the number of true positives, true negatives, false positives and false negatives, respectively. All metrics were implemented using functions from the metrics module in scikit-learn, including accuracy_score, precision_score, recall_score, f1_score, roc_curve, and auc.

### FluRisk web server implementation

The FluRisk web platform was developed using Tornado as the core Python web framework and asynchronous networking library, complemented by Celery for distributed task processing and message handling to enable asynchronous notifications and computational job management, while client-server communication was implemented through JavaScript Object Notation (JSON) and dynamic interface components were created using JQuery and Bootstrap frameworks for interactive data visualization, with additional integration of the DataTables library for enhanced table functionality throughout the web application.

### Data availability statement

All research data supporting this study, including the complete molecular marker atlas, genomic sequences utilized for Flu Host Predictor (FHP) and reference strain analyses, along with mammalian virulence and receptor-binding datasets employed in Flu Virulence Predictor (FVP) and Flu Receptor-binding preference Predictor (FRP), are publicly accessible through the FluRisk web server platform.

## Acknowledgements

We thank members in PengLab for helpful discussions on the manuscript. This work was supported by National Key Plan for Scientific Research and Development of China (2022YFC2303802), the R&D Program of Guangzhou National Laboratory (GZNL2024A01002), National Natural Science Foundation of China (32170651 & 32370700), and Hunan Provincial Natural Science Foundation of China (2024JJ2015).

## References

1. Krammer, F. et al. Influenza. Nat Rev Dis Primers 4, 3 (2018).

2. Short, K. R. et al. One health, multiple challenges: The inter-species transmission of influenza A virus. One Health 1, 1–13 (2015).

3. Du, R., Cui, Q. & Rong, L. Flu Universal Vaccines: New Tricks on an Old Virus. VS 36, 13–24 (2021).

4. Subbarao, K. Avian influenza H7N9 viruses: a rare second warning. Cell Res 28, 1–2 (2018).

5. Yang, L. et al. Genesis and Spread of Newly Emerged Highly Pathogenic H7N9 Avian Viruses in Mainland China. Journal of Virology 91, e01277–17 (2017).

6. Bi, Y. et al. Highly pathogenic avian influenza H5N1 Clade 2.3.2.1c virus in migratory birds, 2014-2015. VS 31, 300–305 (2016).

7. Peacock, T. P. et al. The global H5N1 influenza panzootic in mammals. Nature 637, 304–313 (2025).

8. Puryear, W. B. & Runstadler, J. A. High-pathogenicity avian influenza in wildlife: a changing disease dynamic that is expanding in wild birds and having an increasing impact on a growing number of mammals. javma 262, 601–609 (2024).

9. Sah, R. et al. Concerns on H5N1 avian influenza given the outbreak in U.S. dairy cattle. The Lancet Regional Health - Americas 35, 100785 (2024).

10. Nidom, C. A. et al. Influenza A (H5N1) Viruses from Pigs, Indonesia. Emerg Infect Dis 16, 1515–1523 (2010).

11. Domańska-Blicharz, K. et al. Outbreak of highly pathogenic avian influenza A(H5N1) clade 2.3.4.4b virus in cats, Poland, June to July 2023. Euro Surveill 28, 2300366 (2023).

12. Sevilla, N. et al. Highly pathogenic avian influenza A (H5N1) virus outbreak in Peru in 2022–2023. Infectious Medicine 3, 100108 (2024).

13. EFSA. FLURISK project proposes a model to rank animal influenza strains by their potential to infect humans. https://www.ecdc.europa.eu/en/news-events/flurisk-project-proposes-model-rank-animal-influenza-strains-their-potential-infect (2014).

14. CDC. Influenza Risk Assessment Tool (IRAT) Virus Descriptions and Report Summaries. *Pandemic Flu* https://www.cdc.gov/pandemic-flu/php/monitoring/virus-description.html (2025).

15. WHO. Tool for Influenza Pandemic Risk Assessment (TIPRA). https://www.who.int/teams/global-influenza-programme/avian-influenza/tool-for-influenza-pandemic-risk-assessment-(tipra) (2016).

16. Shu, Y. & McCauley, J. GISAID: Global initiative on sharing all influenza data – from vision to reality. Euro Surveill 22, 30494 (2017).

17. Peng, Y. et al. Automated recommendation of the seasonal influenza vaccine strain with PREDAC. Biosafety and Health 02, 117–119 (2020).

18. Yin, R. et al. ViPal: A framework for virulence prediction of influenza viruses with prior viral knowledge using genomic sequences. Journal of Biomedical Informatics 142, 104388 (2023).

19. Mock, F., Viehweger, A., Barth, E. & Marz, M. VIDHOP, viral host prediction with deep learning. Bioinformatics 37, 318–325 (2021).

20. Borkenhagen, L. K., Allen, M. W. & Runstadler, J. A. Influenza virus genotype to phenotype predictions through machine learning: a systematic review. Emerg Microbes Infect 10, 1896–1907 (2021).

21. Mok, C. K. P. et al. Amino Acid Substitutions in Polymerase Basic Protein 2 Gene Contribute to the Pathogenicity of the Novel A/H7N9 Influenza Virus in Mammalian Hosts. Journal of Virology 88, 3568–3576 (2014).

22. Zhang, H. et al. The PB2 E627K mutation contributes to the high polymerase activity and enhanced replication of H7N9 influenza virus. Journal of General Virology 95, 779–786 (2014).

23. Xu, G. et al. Prevailing PA Mutation K356R in Avian Influenza H9N2 Virus Increases Mammalian Replication and Pathogenicity. Journal of Virology 90, 8105–8114 (2016).

24. Zhang, X. et al. PB1 S524G mutation of wild bird-origin H3N8 influenza A virus enhances virulence and fitness for transmission in mammals. Emerging Microbes & Infections 10, 1038–1051 (2021).

25. Lin, T.-H. et al. A single mutation in bovine influenza H5N1 hemagglutinin switches specificity to human receptors. Science 386, 1128–1134 (2024).

26. Bloom, J. D., Gong, L. I. & Baltimore, D. Permissive Secondary Mutations Enable the Evolution of Influenza Oseltamivir Resistance. Science 328, 1272–1275 (2010).

27. Zhou, L. et al. A single N342D substitution in Influenza B Virus NA protein determines viral pathogenicity in mice. Emerging Microbes & Infections (2020).

28. Lu, C. et al. FluPhenotype—a one-stop platform for early warnings of the influenza A virus. Bioinformatics 36, 3251–3253 (2020).

29. Yin, R., Luo, Z., Zhuang, P., Lin, Z. & Kwoh, C. K. VirPreNet: a weighted ensemble convolutional neural network for the virulence prediction of influenza A virus using all eight segments. Bioinformatics 37, 737–743 (2021).

30. Schrauwen, E. J. & Fouchier, R. A. Host adaptation and transmission of influenza A viruses in mammals. Emerg Microbes Infect 3, 1–10 (2014).

31. Long, J. S., Mistry, B., Haslam, S. M. & Barclay, W. S. Host and viral determinants of influenza A virus species specificity. Nat Rev Microbiol 17, 67–81 (2019).

32. Hie, B., Zhong, E. D., Berger, B. & Bryson, B. Learning the language of viral evolution and escape. Science 371, 284–288 (2021).

33. Wang, G. et al. Deep-learning-enabled protein–protein interaction analysis for prediction of SARS-CoV-2 infectivity and variant evolution. Nat Med 29, 2007–2018 (2023).

34. Nie, Z. et al. A unified evolution-driven deep learning framework for virus variation driver prediction. Nat Mach Intell 7, 131–144 (2025).

35. Fu, L., Niu, B., Zhu, Z., Wu, S. & Li, W. CD-HIT: accelerated for clustering the next-generation sequencing data. Bioinformatics 28, 3150–3152 (2012).

36. Peng, Y. et al. Identification of genome-wide nucleotide sites associated with mammalian virulence in influenza A viruses. Biosafety and Health 2, 32–38 (2020).

37. Raman, R. et al. Advancing glycomics: Implementation strategies at the Consortium for Functional Glycomics. Glycobiology 16, 82R–90R (2006).

38. Suenaga, E., Mizuno, H. & Kumar, P. K. R. Influenza virus surveillance using surface plasmon resonance. Virulence 3, 464–470 (2012).

39. Verzijl, D., Riedl, T., Parren, P. W. H. I. & Gerritsen, A. F. A novel label-free cell-based assay technology using biolayer interferometry. Biosensors and Bioelectronics 87, 388–395 (2017).

40. Aydin, S. A short history, principles, and types of ELISA, and our laboratory experience with peptide/protein analyses using ELISA. Peptides 72, 4–15 (2015).

41. Pedregosa, F. et al. Scikit-learn: Machine Learning in Python. J. Mach. Learn. Res. 12, 2825–2830 (2011).

42. Genuer, R., Poggi, J.-M. & Tuleau-Malot, C. VSURF: An R Package for Variable Selection Using Random Forests. The R Journal 7, 19–33 (2015).

43. Camacho, C. et al. BLAST+: architecture and applications. BMC Bioinformatics 10, 1–9 (2009).

