## Supplementary Materials for "Risk evaluation of newly emerging flu viruses based on genomic sequences and AI"

**Figure S1** The workflow of predicting influenza virus host, mammalian virulence and receptor-binding preference using machine-learning methods.

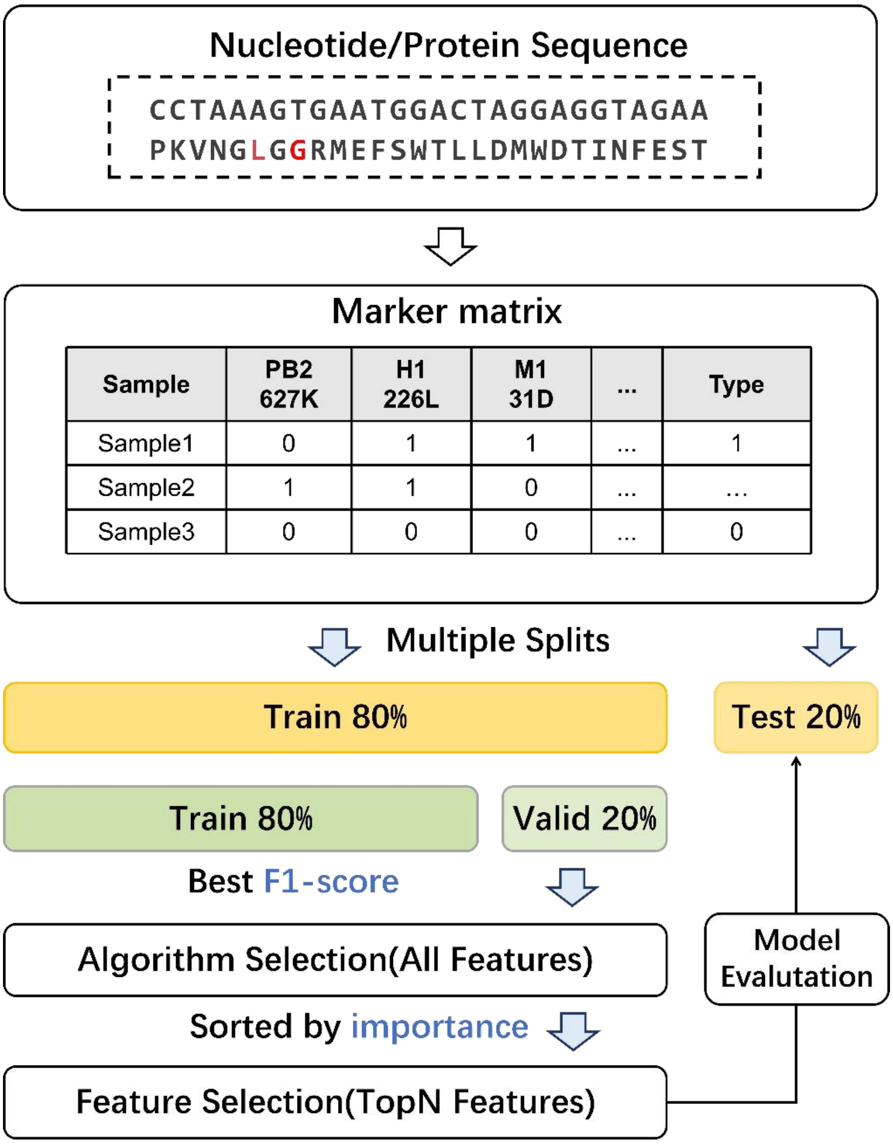


**Figure S2** Estimation of drug-resistance of influenza viruses based on molecular markers using a reference-based method. (A)-(C) shows the distribution of the number of molecular markers and the cumulative distribution of the number of molecular markers associated with laninamivir, baloxavir, and amantadine, respectively.

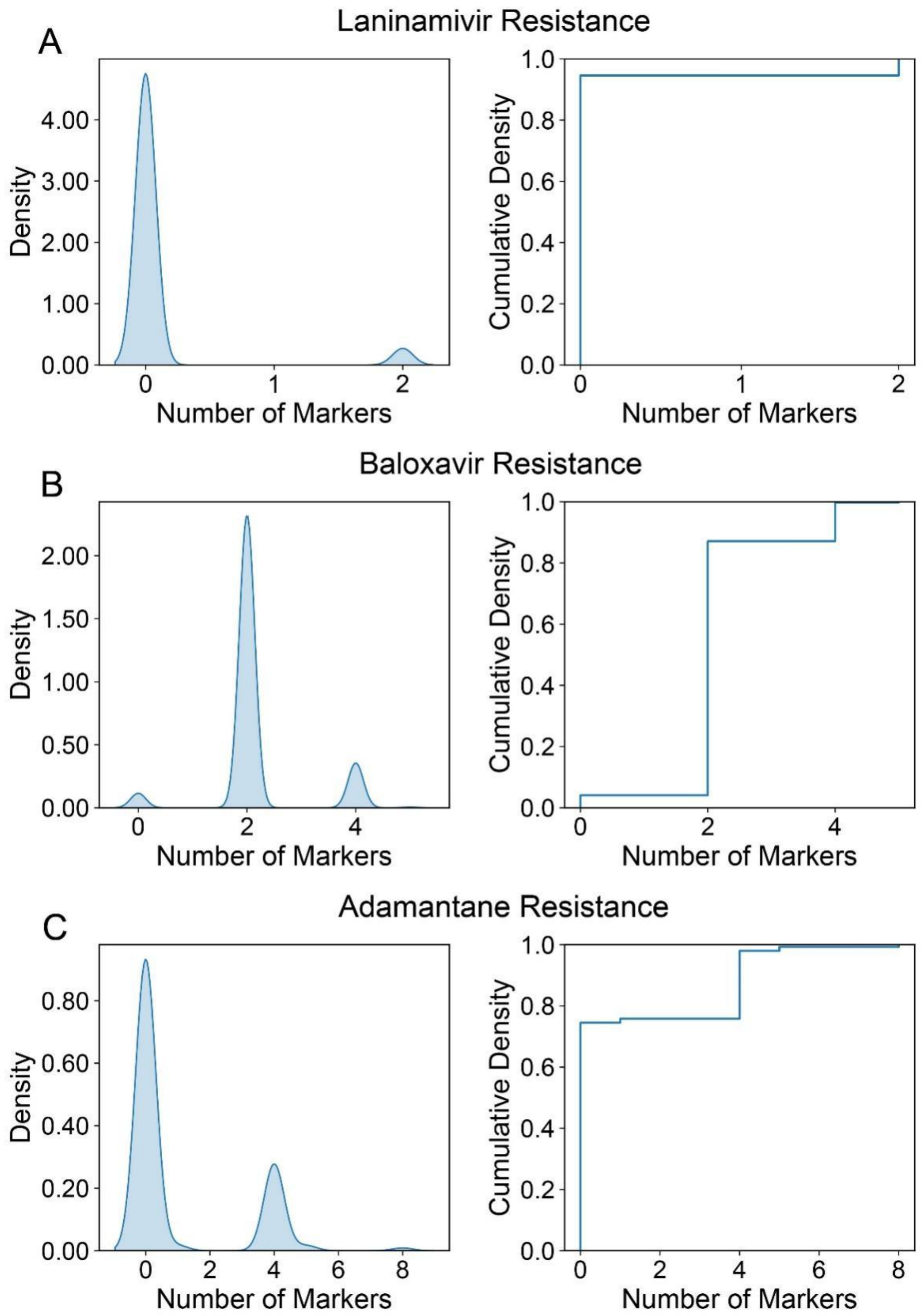

Laninamivir Resistance
A
Density
4.00
3.00
2.00
1.00
0.00
0
1
2
Number of Markers
Cumulative Density
1.0
0.8
0.6
0.4
0.2
0.0
Number of Markers
Baloxavir Resistance
B
Density
2.00
1.50
1.00
0.50
0.00
0
2
4
Number of Markers
Cumulative Density
Number of Markers
Adamantane Resistance
C
Density
0.80
0.60
0.40
0.20
0.00
0
2
4
6
8
Number of Markers
Cumulative Density
Number of Markers

**Table S1**. Comparison of FVP and two previous methods in predicting mammalian virulence. The model performances of two previous methods (ViPal and VirPreNet) were directed compiled from their papers[1,2]. FVP was trained and tested using the same dataset and strategy as these methods. Values represent mean (standard deviation).

| Method* | Accuracy | Precision | Recall | F1-score | AUC |
|---|---|---|---|---|---|
| ViPal | 0.802(0.041) | 0.801(0.040) | 0.879(0.074) | 0.842(0.035) | 0.749(0.082) |
| VirPreNet | 0.794(0.028) | 0.781(0.044) | **0.948**(0.029) | 0.856(0.025) | - |
| FVP | **0.811**(0.030) | **0.816**(0.036) | 0.945(0.031) | **0.875**(0.019) | **0.781**(0.059) |

**References**

1 Yin, R. *et al.* ViPal: A framework for virulence prediction of influenza viruses with prior viral knowledge using genomic sequences. *Journal of Biomedical* 586 *Informatics* **142**, 104388 (2023)

2 Yin, R., Luo, Z., Zhuang, P., Lin, Z. & Kwoh, C. K. VirPreNet: a weighted ensemble convolutional neural network for the virulence prediction of influenza A virus using all eight segments. *Bioinformatics* **37**, 737–743 (2021).
